# ERGA-BGE Reference Genome of the Striped Field Mouse (*Apodemus agrarius*), a Widespread and Abundant Species in Central and Eastern Europe

**DOI:** 10.1101/2024.12.04.626796

**Authors:** Franc Janžekovič, Elena Buzan, Aja Bončina, Nuria Escudero, Rosa Fernández, Astrid Böhne, Rita Monteiro, Laura Aguilera, Marta Gut, Francisco Câmara Ferreira, Fernando Cruz, Jèssica Gómez-Garrido, Tyler S. Alioto, Leanne Haggerty, Fergal Martin, Diego De Panis

## Abstract

The reference genome of *Apodemus agrarius* provides a valuable resource for phylogenetic studies of rodents, particularly mice, and for understanding factors that influence the geographical distribution of the species across East Asia and East Europe. A total of 25 contiguous chromosomal pseudomolecules were assembled from the genome sequence (23 autosomes and 2 sex chromosomes). This chromosome-level assembly encompasses 2.6 Gb, composed of 242 contigs and 60 scaffolds, with contig and scaffold N50 values of 35 Mb and 119 Mb, respectively.

## Introduction

The genus *Apodemus* belongs to the order Rodentia, family Muridae. The striped field mouse is classified in the subgenus *Apodemus* (sensu stricto) (Wilson et al., 2017). This species has an extensive yet disjunct distribution, spanning two major regions: the Palearctic in the west and the Indomalayan in the east. The western range includes central and eastern Europe, extending across the Caucasus, while the eastern range covers areas of East Asia (Gliwicz, J. and Kryštufek, 1999; Wilson et al., 2017).

The striped field mouse is a diurnal species found in a range of terrestrial habitats including woodland edges, grasslands, marshes, reedbeds, cornfields, pastures, gardens in rural and suburban areas, and green spaces in urban areas. Moist habitats are preferred. It feeds on roots, grains, seeds, berries, nuts, and insects. Additionally, it is a crucial food source for birds of prey, such as owls, as well as smaller carnivores (Wilson et al., 2017).

The generation of this reference resource was coordinated by the European Reference Genome Atlas (ERGA) initiative’s Biodiversity Genomics Europe (BGE) project, supporting ERGA’s aims of promoting transnational cooperation to promote advances in the application of genomics technologies to protect and restore biodiversity (Mazzoni et al., 2023).

## Materials & Methods

ERGA’s sequencing strategy includes Oxford Nanopore Technology (ONT) and/or Pacific Biosciences (PacBio) for long-read sequencing, along with Hi-C sequencing for chromosomal architecture, Illumina Paired-End (PE) for polishing (i.e. recommended for ONT-only assemblies), and RNA sequencing for transcriptomic profiling, to facilitate genome assembly and annotation.

### Sample and Sampling Information

On October 20^th^ 2022, two adult male *Apodemus agrarius* specimens were collected by Franc Janžekovič in Dravsko Polje, Slovenia. The specimens were morphologically identified using illustrated (Wilson et al., 2017) and dichotomous (Kryštufek & Janžekovič, 1999) keys. The specimen was collected using Sherman traps. No sampling permits were required. Tissue samples (brain, kidney, liver, muscle, testis) were snap-frozen immediately after harvesting and stored in liquid nitrogen until DNA/RNA extraction.

### Vouchering information

Physical reference materials for the sequenced specimen have been deposited in the Slovenian Museum of Natural History, mammal collection (zbirka sesalcev) under the accession numbers PMSL31475 and PMSL31476.

Frozen reference tissue material from muscle is available from the same individuals at the Biobank Slovenian Museum of Natural History under the vouchers PMS_TIS1 and PMS_TIS2.

### Data Availability

*A. agrarius* and the related genomic study were assigned to Tree of Life ID (ToLID) mApoAgr2 and all sample, sequence, and assembly information are available under the umbrella BioProject PRJEB74497. The sample information is available at the following BioSample accessions: SAMEA112797470, SAMEA112797471, and SAMEA112797479. The genome assembly is accessible from ENA under accession number GCA_964023405.1 and the annotated genome is available through the Ensembl Rapid Release page (projects.ensembl.org/erga-bge). Sequencing data produced as part of this project are available from ENA at the following accessions: ERX12202498, ERX13168343, ERX13168342, and ERX12202500. Documentation related to the genome assembly and curation can be found in the ERGA Assembly Report (EAR) document available at github.com/ERGA-consortium/EARs/tree/main/Assembly_Reports/Apodemus_agrarius/mApoAgr2.

Further details and data about the project are hosted on the ERGA portal at portal.erga-biodiversity.eu/data_portal/39030.

### Genetic Information

The estimated genome size for the subfamily Murinae, based on several species, is 3.11 Gb, while the estimation based on reads kmer profiling is 2.82 Gb. This is a diploid genome with a haploid number of 24 chromosomes (2n=48), including XY sex chromosomes in males. All information for this species was retrieved from Genomes on a Tree (Challis et al., 2023).

### DNA/RNA processing

DNA was extracted from kidney tissue using the Blood & Cell Culture DNA Midi Kit (Qiagen) following the manufacturer’s instructions. DNA quantification was performed using a Qubit dsDNA BR Assay Kit (Thermo Fisher Scientific), and DNA integrity was assessed using a Femtopulse system (Genomic DNA 165 Kb Kit, Agilent). DNA was stored at 4ºC until use.

RNA was extracted from testis tissue using an RNeasy Mini Kit (Qiagen) according to the manufacturer’s instructions. RNA quantification was performed using the Qubit RNA BR Kit and RNA integrity was assessed using a Bioanalyzer 2100 system (Eukaryote Total RNA Pico Kit, Agilent). RNA was stored at -80ºC until use.

### Library Preparation and Sequencing

A long-read whole genome library was prepared using the SQK-LSK114 kit and sequenced on a PromethION P24 A series instrument (Oxford Nanopore Technologies). For short-read whole genome sequencing (WGS), a library was prepared using the KAPA Hyper Prep Kit (Roche). A Hi-C library was prepared from kidney tissue using the Dovetail Omni-C Kit (CantataBio), followed by the KAPA Hyper Prep Kit for Illumina sequencing (Roche). The RNA library, generated from testis tissue, was prepared with the KAPA mRNA Hyper Prep Kit (Roche). All the short-read libraries were sequenced on the Illumina NovaSeq 6000 instrument. In total, 91x Oxford Nanopore, 87x Illumina WGS shotgun, and 64x HiC data were sequenced to generate the assembly.

### Genome Assembly Methods

The genome was assembled using the CNAG CLAWS pipeline (Gomez-Garrido, 2024). Briefly, reads were preprocessed for quality and length using Trim Galore v0.6.7 and Filtlong v0.2.1, and initial contigs were assembled using NextDenovo v2.5.0, followed by polishing of the assembled contigs using HyPo v1.0.3, removal of retained haplotigs using purge-dups v1.2.6 and scaffolding with YaHS v1.2a. Finally, assembled scaffolds were curated via manual inspection using Pretext v0.2.5 with the Rapid Curation Toolkit (gitlab.com/wtsi-grit/rapid-curation) to remove any false joins and incorporate any sequences not automatically scaffolded into their respective locations in the chromosomal pseudomolecules (or super-scaffolds). Finally, the mitochondrial genome was assembled as a single circular contig of 16,433 bp using the FOAM pipeline v0.5 (github.com/cnag-aat/FOAM) and included in the released assembly (GCA_963981305.1). Summary analysis of the released assembly was performed using the ERGA-BGE Genome Report ASM Galaxy workflow (De Panis, 2024b), incorporating tools such as BUSCO v5.5, Merqury v1.3, and others (see reference for the full list of tools).

### Genome Annotation Methods

A gene set was generated using the Ensembl Gene Annotation system (Aken et al., 2016), primarily by aligning publicly available short-read RNA-seq data from BioSample SAMEA112797471 to the genome. Gaps in the annotation were filled via protein-to-genome alignments of a select set of vertebrate proteins from UniProt (The UniProt Consortium, 2019), which had experimental evidence at the protein or transcript level. At each locus, data were aggregated and consolidated, prioritising models derived from RNA-seq data, resulting in a final set of gene models and associated non-redundant transcript sets. To distinguish true isoforms from fragments, the likelihood of each open reading frame (ORF) was evaluated against known vertebrate proteins. Low-quality transcript models, such as those showing evidence of fragmented ORFs, were removed. In cases where RNA-seq data were fragmented or absent, homology data were prioritised, favouring longer transcripts with strong intron support from short-read data. The resulting gene models were classified into three categories: protein-coding, pseudogene, and long non-coding. Models with hits to known proteins and few structural abnormalities were classified as protein-coding. Models with hits to known proteins but displaying abnormalities, such as the absence of a start codon, non-canonical splicing, unusually small intron structures (<75 bp), or excessive repeat coverage, were reclassified as pseudogenes. Single-exon models with a corresponding multi-exon copy elsewhere in the genome were classified as processed (retrotransposed) pseudogenes. Models that did not fit any of the previously described categories did not overlap protein-coding genes, and were constructed from transcriptomic data were considered potential lncRNAs. Potential lncRNAs were further filtered to remove single-exon loci due to their unreliability. Putative miRNAs were predicted by performing a BLAST search of miRBase (Kozomara et al., 2019) against the genome, followed by RNAfold analysis (Gruber et al., 2008). Other small non-coding loci were identified by scanning the genome with Rfam (Kalvari et al., 2018) and passing the results through Infernal (Nawrocki & Eddy, 2013). Summary analysis of the released annotation was performed using the ERGA-BGE Genome Report ANNOT Galaxy workflow (De Panis, 2024a) (De Panis, 2024), incorporating tools such as AGAT v1.2, OMArk v0.3, and others (see reference for the full list of tools).

## Results

### Genome Assembly

The genome assembly has a total length of 2,613,539,280 bp in 61 scaffolds including the mitogenome (Figures 1 and 2), with a GC content of 42.15%. It features a contig N50 of 35,188,814 bp (L50=22) and a scaffold N50 of 119,125,257 bp (L50=9). There are 182 gaps, totaling 36,400 kb in cumulative size. The single-copy gene content analysis using the Mammalia database with BUSCO resulted in 96.8% completeness (95.5% single and 1.3% duplicated). 90.9% of reads k-mers were present in the assembly and the assembly has a base accuracy Quality Value (QV) of 47.1 as calculated by Merqury.

**Figure 1.**
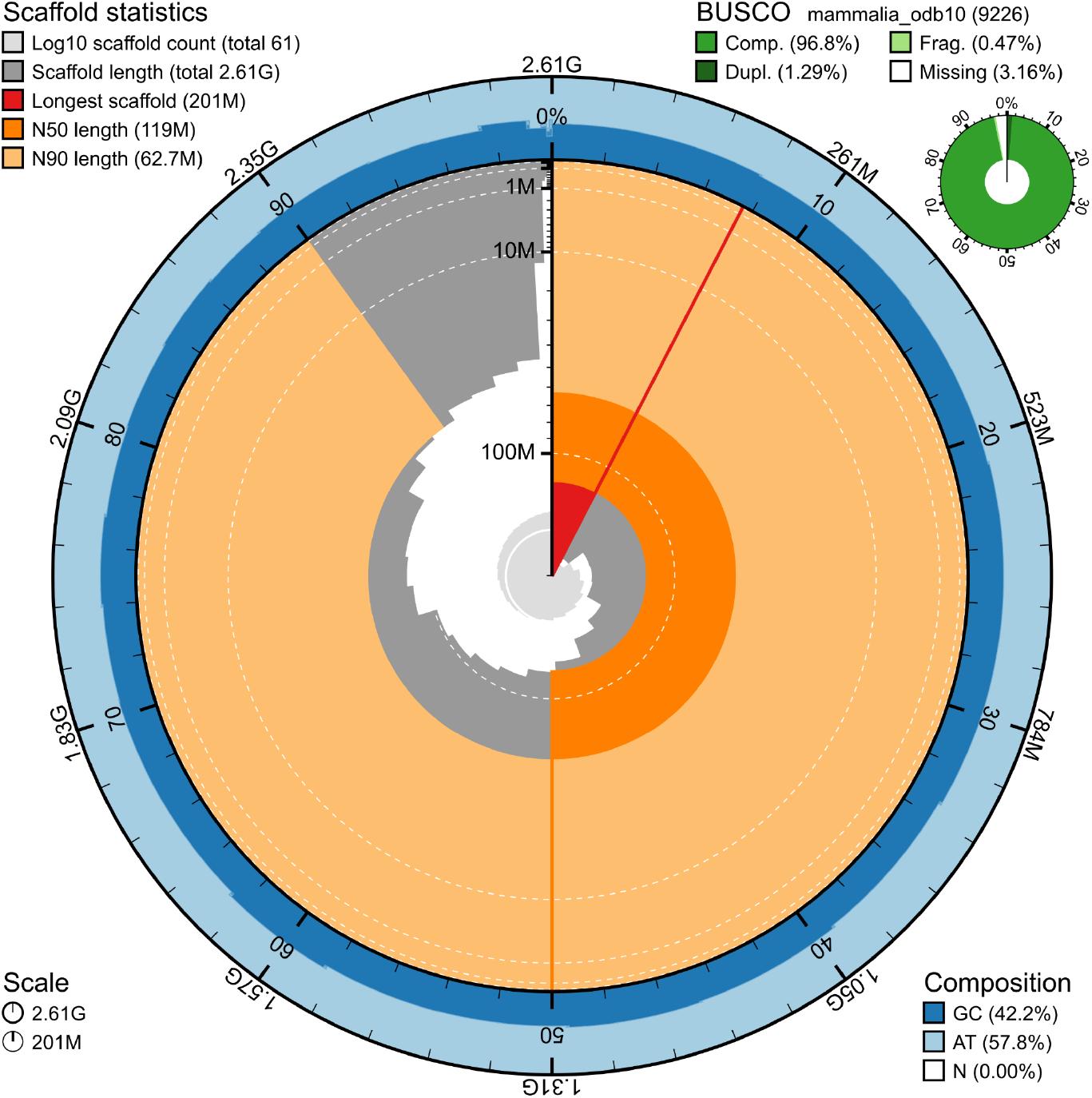
Snail plot summary of assembly statistics. The main plot is divided into 1,000 size-ordered bins around the circumference, with each bin representing 0.1% of the 2,613,539,280 bp assembly including the mitochondrial genome. The distribution of sequence lengths is shown in dark grey, with the plot radius scaled to the longest sequence present in the assembly (200,613,070 bp, shown in red). Orange and pale-orange arcs show the scaffold N50 and N90 sequence lengths (119,125,257 and 62,738,681 bp), respectively. The pale grey spiral shows the cumulative sequence count on a log-scale, with white scale lines showing successive orders of magnitude. The blue and pale-blue area around the outside of the plot shows the distribution of GC, AT, and N percentages in the same bins as the inner plot. A summary of complete, fragmented, duplicated, and missing BUSCO genes found in the assembled genome from the Mammalia database (odb10) is shown on the top right.

**Figure 2.**
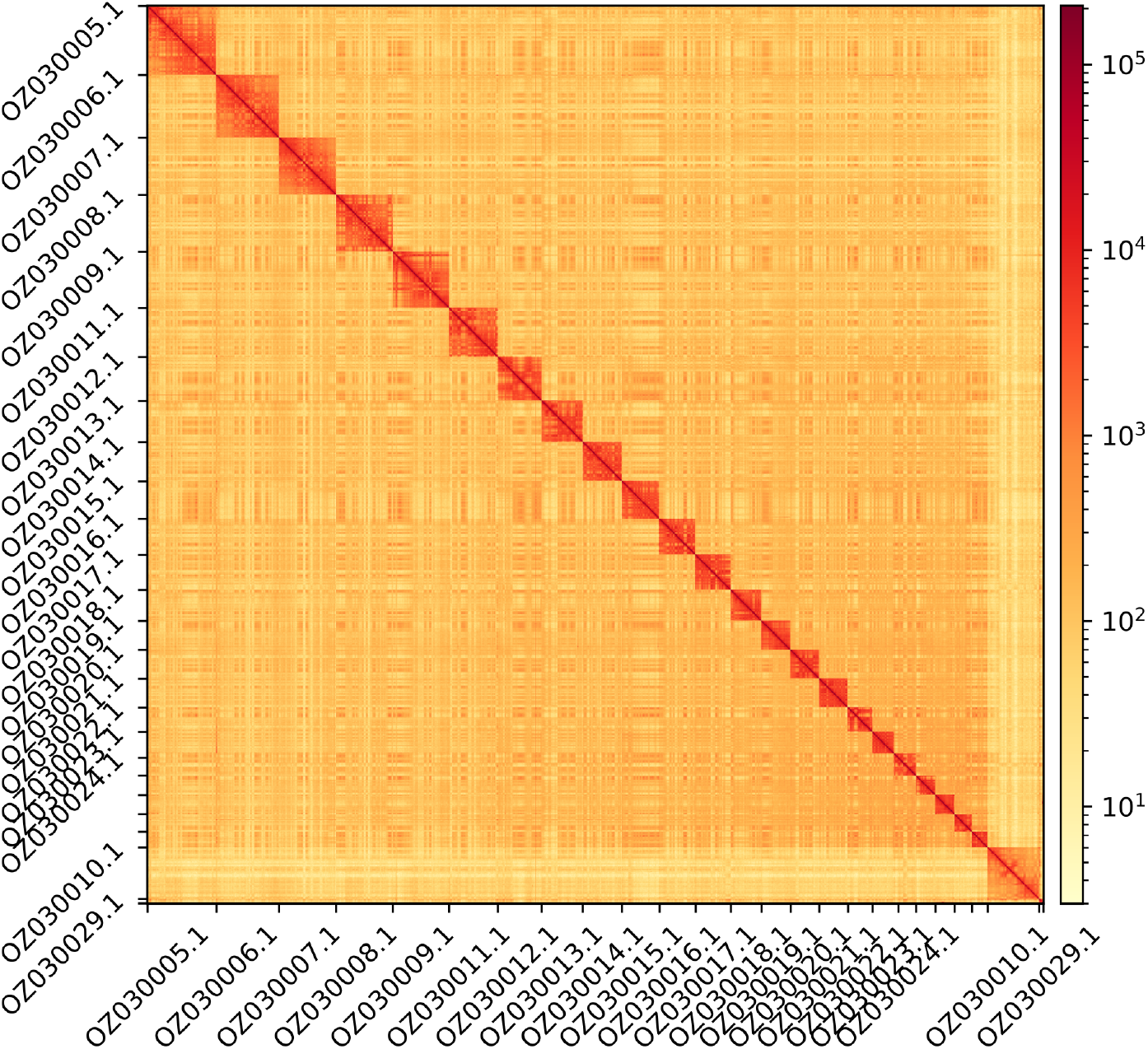
Hi-C contact map showing spatial interactions between regions of the genome. The diagonal corresponds to intra-chromosomal contacts, depicting chromosome boundaries. The frequency of contacts is shown on a logarithmic heatmap scale. Hi-C matrix bins were merged into a 50 kb bin size for plotting. Due to space constraints on the axes, only the GenBank names of the 19th largest autosomes and the sex chromosomes are shown.

### Genome Annotation

The genome annotation consists of 20,679 protein-coding genes with an associated 28,164 transcripts, in addition to 260 immune system receptor segment genes, 485 pseudogenes, and 3,956 non-coding RNA genes of various types (Table 1). Using the longest isoform per transcript, the single-copy gene content analysis using the Mammalia database with BUSCO resulted in 98.8% completeness. Using the OMAmer Myomorpha database for OMArk resulted in 96.14% completeness and 98.40% consistency (Table 2).

**Table 1.** Statistics from assembled gene models.

| Category | No. genes | No. transcripts | Mean* gene length (bp) | No. single-exon genes | Mean* exons per transcript |
| --- | --- | --- | --- | --- | --- |
| Protein-coding | 20,679 | 28,164 | 42,555 | 2,662 | 10.6 |
| Ig/TCR segments | 260 | 260 | 78-5,910 | 170 | 1.0-1.4 |
| Pseudogenes | 485 | 485 | 5,402 | 34 | 6.6 |
| lncRNA | 352 | 440 | 18,266 | 67 | 2.6 |
| snRNA | 1,599 | 1,599 | 117 | 1,599 | 1.0 |
| snoRNA | 1,555 | 1,555 | 119 | 1,555 | 1.0 |
| rRNA | 63 | 63 | 314 | 63 | 1.0 |
| tRNA | 22 | 22 | 68 | 22 | 1.0 |
| miRNA | 87 | 87 | 87 | 87 | 1.0 |
| scRNA | 32 | 32 | 169 | 32 | 1.0 |
| Other non-coding | 246 | 246 | 108-275 | 246 | 1.0-1.0 |

\*The range of the mean values is shown in combined categories

**Table 2.** Annotation completeness and consistency scores calculated by BUSCO run in protein mode (Mammalia) and OMArk (Myomorpha)

|  | Complete | Singular | Duplicated | Fragmented | Missing |
| --- | --- | --- | --- | --- | --- |
| BUSCO | 9,118 (98.8%) | 9,055 (98.1%) | 63 (0.7%) | 27 (0.3%) | 81 (0.9%) |
| OMArk | 15,662 (98.50%) | 15,286 (96.14%) | 376 (2.36%) | - | 237 (1.49%) |
|  | Consistent | Inconsistent | Contaminants | Unknown |  |
| OMArk | 20,604 (98.40%) | 237 (1.13%) | 0 (0.00%) | 98 (0.47%) |  |

## Acknowledgements

We want to express our gratitude to researcher Janko Skok from the University of Maribor for their invaluable assistance in field sampling and to researcher Aja Bončina from the University of Primorska for her help with the conservation of the samples. We acknowledge the support of the Freiburg Galaxy Team: Saim Momin and Björn Grüning, Bioinformatics, University of Freiburg (Germany), funded by the German Federal Ministry of Education and Research BMBF grant 031 A538A de.NBI-RBC and the Ministry of Science, Research and the Arts Baden-Württemberg (MWK) within the framework of LIBIS/de.NBI Freiburg. We would like to acknowledge the assembly reviewer, Joanna Collins from the Wellcome Sanger Institute.

## Conflict of Interest

The authors declare no conflict of interest related to this study. The funding sources had no involvement in the study design, collection, analysis, or interpretation of data; in the writing of the manuscript; or in the decision to submit the article for publication. All authors have participated sufficiently in the work to take public responsibility for the content and agree to the submission of this manuscript.

## Funder Information

Biodiversity Genomics Europe **(Grant no.101059492)** is funded by Horizon Europe under the Biodiversity, Circular Economy and Environment call (REA.B.3); co-funded by the Swiss State Secretariat for Education, Research and Innovation (SERI) under contract numbers 22.00173 and 24.00054; and by the UK Research and Innovation (UKRI) under the Department for Business, Energy and Industrial Strategy’s Horizon Europe Guarantee Scheme. The Slovenian Research and Innovation Agency supported the research of FJ with the Research Programme (P1-0403).

## Author Contributions

FJ and EB coordinated the project, FJ collected the species, FJ, AB, and EB identified the species, preserved biological material, and provided metadata, NE, RF, RM, and AB provided sampling and metadata support and management, LA and MG extracted DNA, prepared libraries, and performed sequencing, FCF, JGG and FC performed genome assembly and curation under the supervision of TSA, DDP generated the analysis and report. All authors contributed to the writing, review, and editing of this genome note and read and approved the final version.

## Notes

### Competing Interest Statement

The authors have declared no competing interest.

## Literature Cited

Aken, B. L., Ayling, S., Barrell, D., Clarke, L., Curwen, V., Fairley, S., Fernandez Banet, J., Billis, K., García Girón, C., Hourlier, T., Howe, K., Kähäri, A., Kokocinski, F., Martin, F. J., Murphy, D. N., Nag, R., Ruffier, M., Schuster, M., Tang, Y. A., … Searle, S. M. J. (2016). The Ensembl gene annotation system. Database, 2016, baw093. 10.1093/database/baw093

Challis, R., Kumar, S., Sotero-Caio, C., Brown, M., & Blaxter, M. (2023). Genomes on a Tree (GoaT): A versatile, scalable search engine for genomic and sequencing project metadata across the eukaryotic tree of life. Wellcome Open Research, 8, 24. 10.12688/wellcomeopenres.18658.1

De Panis, D. (2024a). ERGA-BGE Genome Report ANNOT analyses. WorkflowHub. 10.48546/WORKFLOWHUB.WORKFLOW.1096.1

De Panis, D. (2024b). ERGA-BGE Genome Report ASM analyses (one-asm WGS Illumina PE + HiC). WorkflowHub. 10.48546/WORKFLOWHUB.WORKFLOW.1103.2

Gomez-Garrido, J. (2024). CLAWS (CNAG’s long-read assembly workflow in Snakemake). WorkflowHub. 10.48546/WORKFLOWHUB.WORKFLOW.567.2

Gruber, A. R., Lorenz, R., Bernhart, S. H., Neuböck, R., & Hofacker, I. L. (2008). The Vienna RNA Websuite. Nucleic Acids Research, 36(Suppl_2), W70–W74. 10.1093/nar/gkn188

Kalvari, I., Nawrocki, E. P., Argasinska, J., Quinones-Olvera, N., Finn, R. D., Bateman, A., & Petrov, A. I. (2018). Non-Coding RNA Analysis Using the Rfam Database. Current Protocols in Bioinformatics, 62(1), e51. 10.1002/cpbi.51

Kozomara, A., Birgaoanu, M., & Griffiths-Jones, S. (2019). miRBase: From microRNA sequences to function. Nucleic Acids Research, 47(D1), D155–D162. 10.1093/nar/gky1141

Kryštufek, B., & Janžekovič, F. (1999). Ključ za določanje vretenčarjev Slovenije. DZS. https://www.dlib.si/details/URN:NBN:SI:DOC-TDK1RUJ4/?query=%27contributor%3DVrezec%2C+Al%27&pageSize=25&fUDC=Biologija

Mazzoni, C., Ciofi, C., & Waterhouse, R. (2023). Biodiversity: An atlas of European reference genomes. Nature, 619, 252–252. 10.1038/d41586-023-02229-w

Mitchell-Jones, A. J., Amori, G., Bogdanowicz, W., Krystufek, B., Reijnders, P. J. H., Spitzenberger, F., Stubbe, M., Thissen, J. B. M., Vohralik, V., & Zima, J. (1999). The atlas of European mammals. Poyser. https://research.wur.nl/en/publications/the-atlas-of-european-mammals

Nawrocki, E. P., & Eddy, S. R. (2013). Infernal 1.1: 100-fold faster RNA homology searches. Bioinformatics, 29(22), 2933–2935. 10.1093/bioinformatics/btt509

The UniProt Consortium. (2019). UniProt: A worldwide hub of protein knowledge. Nucleic Acids Research, 47(D1), D506–D515. 10.1093/nar/gky1049

Wilson, D. E., Mittermeier, R. A., & Lacher, T. E. (2017). Handbook of the mammals of the world: Vol. 7 : rodents II (Vol. 7). Lynx Edicions. https://portals.iucn.org/library/node/47516

